# Multi-session tDCS paired with passive mobilisation increases thalamo-cortical coupling during command following

**DOI:** 10.1101/2022.11.22.517479

**Authors:** Davide Aloi, Roya Jalali, Sara Calzolari, Melanie Lafanechere, R. Chris Miall, Davinia Fernández-Espejo

## Abstract

**Background:** Therapeutic options for patients with prolonged disorder of consciousness (PDOC) are limited. PDOC patients often exhibit a dissociation between their retained level of (covert) cognitive ability and their (overt) behavioural responses (cognitive-motor dissociation; CMD). This is linked to reduced coupling between thalamus and the primary motor cortex.

**Objective:** To assess whether pairing tDCS with a concurrent passive mobilisation protocol (designed to be feasible in PDOC) can influence thalamo-M1 dynamics and whether these changes are enhanced after multiple stimulation sessions.

**Methods:** We used Dynamic Causal Modelling (DCM) on functional magnetic resonance imaging (fMRI) data from 22 healthy participants to assess tDCS changes on effective connectivity within motor network areas during command-following.

**Results:** We found that a single anodal tDCS session (paired with passive mobilisation of the thumb) decreased self-inhibition in the motor cortex, with five sessions further enhancing this effect. In addition, anodal tDCS increased thalamo-M1 excitation as compared to cathodal stimulation, with the effects maintained after 5 sessions. In turn, cathodal tDCS had opposing effects on these connections after one session but became more similar to anodal after 5.

**Conclusions:** Together, our results suggest that pairing anodal tDCS with passive mobilisation across multiple sessions may facilitate behavioural command-following in PDOC patients with CMD. More broadly, they offer a mechanistic window into the neural underpinnings of the cumulative effects of multi-session tDCS.

## Introduction

Prolonged disorders of consciousness (PDOC), including the vegetative/unresponsive wakefulness state (VS) and the minimally conscious state (MCS) are catastrophic neurological conditions for which an effective therapeutic approach has yet to be found. In recent years, several studies have used transcranial direct current stimulation (tDCS) in an attempt to induce clinical improvement in VS and MCS patients[1]. However, despite some promising results[2–10], the outcome of these trials has been mixed and inconsistent, making it difficult to draw meaningful conclusions about the effectiveness of tDCS in PDOC [1]. Part of this variability corresponds to well-known limitations present in the broad tDCS literature, such as the effect of inter-individual differences in brain anatomy [11], or a lack of standardisation of protocols that leads to heterogeneity of stimulation parameters across studies. In addition, PDOC originates from different aetiologies, resulting in heterogeneous patterns of brain injury across patients [10,12]. However, as we have previously argued [1,13], a key consideration in this patient group refers to the lack of an accurate and precise definition of the neural basis of consciousness [14], which complicates the identification of effective targets for neuromodulation.

In an earlier tDCS study [13], we switched the focus from the consciousness disorder itself to the patients’ (lack of) responsiveness or ability to produce voluntary motor responses. While behavioural responsiveness, and specifically command-following, is the main clinical indicator of awareness in diagnostic examinations of PDOC, it is now well established that it does not accurately reflect the level of cognitive functioning in many patients who are partially (or in some cases fully) aware but lack voluntary motor control [15,16]. This condition is known as cognitive-motor dissociation (CMD) [16] and is associated with reduced thalamo-cortical coupling owing to selective structural impairments in the pathways connecting the thalamus and primary motor cortex (M1) [17,18]. In our previous study, we showed in the healthy brain that a single session of tDCS over M1 delivered at rest can successfully modulate thalamo-cortical dynamics during a subsequent behavioural command-following task [13], suggesting a potential route to improve responsiveness in PDOC. While our earlier results are encouraging, it is well known that tDCS leads to most reliable behavioural effects when paired with a task that can successfully engage the target network [19–22]. In fact, it is thought that tDCS reinforces the pattern of the underlying brain network activity [22–24], and thus, its effects on brain function are dependent on the ongoing neuronal state. Additionally, clinical work has shown that concurrent stimulation and physical therapy help promote motor rehabilitation, as tDCS appears to enhance the response of neural networks to the therapy, optimising plastic changes as a result [25]. Crucially, most PDOC patients (excluding those at the high end of the MCS spectrum) are by definition unable to execute voluntary external responses, and thus unable to take part in a command-following task. For this reason, existing studies in this patient group have delivered tDCS at rest. However, the literature suggests that passive mobilisation may lead to an engagement of the motor areas similar to active movements [26–30] and could thus be an alternative to the use of a command-following task during tDCS. Importantly, PDOC patients typically receive regular passive mobilisation as maintenance therapy for the clinical management of their spasticity [31]. Therefore, a protocol involving passive mobilisation is clinically applicable in PDOC.

In addition, it is well-known that a single session of tDCS may not be sufficient to induce noticeable behavioural changes [32]. In contrast, multiple sessions can lead to more significant improvements, possibly due to a potentiation of the functional connectivity between the regions activated during learning [33]. The PDOC literature is consistent with this, with studies using several stimulation sessions showing more robust clinical improvements [4,7,8].

Therefore, in this study, we used dynamic causal modelling of fMRI data to explore single- and multi-session effects of tDCS over M1 paired with passive mobilisation of the thumb on thalamo-cortical coupling during command-following. We focused on characterising the mechanisms and effects of tDCS on a well-controlled study in healthy individuals, while keeping our protocol and methods translatable to clinical practice in PDOC. We hypothesised that: (a) the combination of neurostimulation and passive mobilisation will enhance thalamo-cortical coupling beyond what we have previously reported when administering stimulation at rest [13], and (b) such enhancement will be greater after multiple stimulation-mobilisation sessions.

## Materials and Methods

### Participants

Twenty-four right-handed healthy volunteers took part in the study (16 women, 8 men; mean age 24 ± 3 years), of whom 22 completed all three weeks of testing and were therefore included in the analyses (16 women, 6 men; mean age 25 ± 4 years). We recruited participants using advertisements across campus as well as the local Research Participation Scheme. All participants completed a pre-screening protocol to ensure eligibility to safely participate in MRI and tDCS experiments. They all reported no history of psychiatric or neurological disorders, no personal or family history of epilepsy, no use of psychoactive drugs, and normal or corrected vision. We instructed participants to come to the sessions well rested and hydrated, and not to consume coffee or alcohol within the 24 hours before each session, as per tDCS safety regulations [34]. The University of Birmingham’s Science, Technology, Engineering, and Mathematics Ethical Review Committee approved the study and all participants gave written informed consent. We compensated participants with £220 or the equivalent in course credits.

### Experimental Procedure

We used a sham- and polarity-controlled, randomised, blind, crossover design. Our protocol involved 3 testing weeks in which participants received either anodal, cathodal, or sham tDCS, in a counterbalanced order (Fig. 1A). Each week included 5 consecutive daily sessions of stimulation of the same polarity (i.e., one polarity per week). There was at least 1 week break between each polarity (mean days: 9 ± 15). Participants were blind to the polarity being used in each session. Before the first testing session, participants provided informed consent and completed the Edinburgh handedness inventory [35]. Each week, we delivered the first and fifth sessions of stimulation in the MRI scanner (we will refer to these as Day1-MRI and Day5-MRI sessions respectively), where participants also performed a command following task before and after the stimulation (this resulted in 12 fMRI runs of the task per participant). We delivered the second, third, and fourth tDCS sessions in a designated cubicle (we will refer to these as Day2-Lab, Day3-Lab, and Day4-Lab sessions). We screened participants to reconfirm MRI safety before each MRI session and to comply with tDCS safety requirements before every stimulation session. After confirming safety, we set up the tDCS electrodes in the MRI control room before each MRI session, or directly in the testing cubicle before each lab session. Once we had completed the tDCS setup, we gave participants an MRI-compatible joystick (FORP-190 932, Current designs INC., PA USA) to record their responses during an fMRI motor command-following task (CF-fMRI) and to deliver the passive mobilisation. For the MRI sessions, we placed the joystick on the participant’s torso and secured their right thumb to the handle using tape. The joystick was connected to the interface through an optical cable. During the lab sessions we secured the joystick on a desk while the participant was sitting on a chair. In the MRI sessions, participants completed the CF-fMRI task before and after tDCS (paired with passive mobilisations). In the lab sessions, participants only received the stimulation/mobilisations but did not have to perform the task. We used a 1200 Hz sampling frequency of x and y positions to record movements during CF-fMRI. To ensure accurate recordings, we calibrated the joystick at the start of each session. We used a Windows 10 Desktop PC and Matlab 2017b to deliver the stimuli and record the motion tracking data. In the MRI sessions, we presented all visual stimuli via a digital system (VPixx PROPIxx) that projected the image onto a mirror fixed to the head coil, with a visual angle of ~10°. We delivered auditory cues using the SOUNDPixx MRI stereo audio system coupled with disposable noise-attenuating (about 29db+) ear tips (MRIaudio INC), and used an intercom system to communicate with participants during the scanning sessions. In the lab sessions we used an Iiyama 24’’ desktop monitor and provided participants with both earplugs and 3M ear defenders (35 dB).

**Fig. 1.**
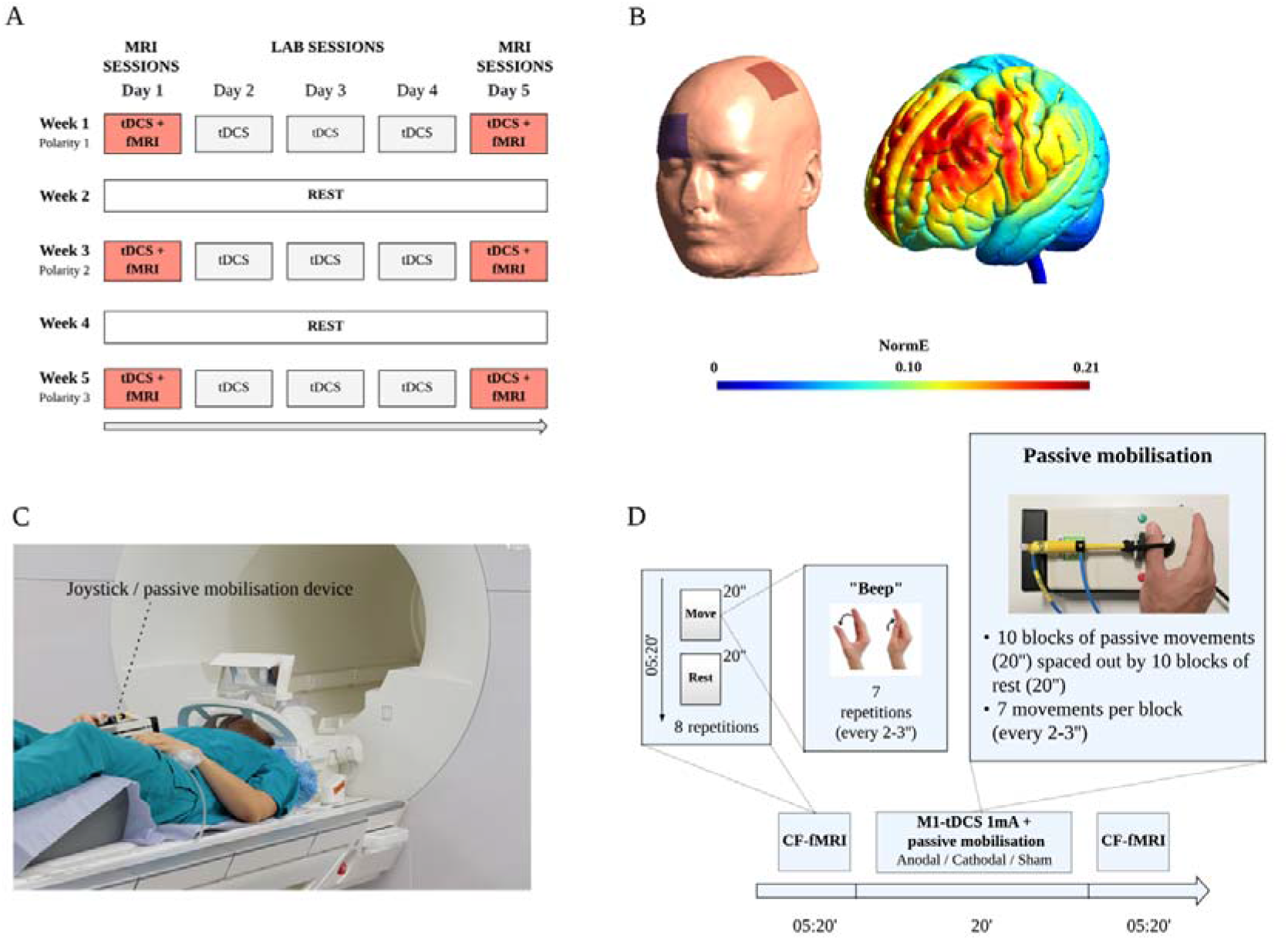
Study Design and tDCS montage. (**A**) Schedule of events for all participants. Note that tDCS was coupled with passive mobilisation of the thumb in all sessions. (**B**) tDCS montage (left) and simulation of current spread (electric field) on the MNI standard head model (right), as calculated with SimNIBS3.2.2. We placed the target electrode on C3 (M1) and the reference electrode on Fp2 for the simulation and set up the current to 1mA. Note that this simulation does not consider inter-individual differences in the position of the electrodes or the different tissue compartments across participants and therefore it should be interpreted as a rough estimate of the canonical field spread that is expected with our montage. (**C**) MRI setup. We recorded participant movements during the command-following (CF-fMRI) task with an MRI compatible joystick, which we also used to deliver passive mobilisation during tDCS. (**D**) MRI session. Participants performed the CF-fMRI task before and after receiving 20 minutes of tDCS coupled with passive mobilisation of the thumb.

After all sessions, participants received a post-tDCS perceptual scale where they rated the sensations and/or discomfort they perceived and indicated whether they thought they received real stimulation or sham.

### Electrical Stimulation

To administer the stimulation, we used a NeuroConn DC-STIMULATOR-MR for the MRI sessions and a DC-STIMULATOR for the lab sessions (neuroCare Group GmbH, Germany). We used 5×5 cm2 electrodes with a thin layer of electro-conductive Ten20® paste to increase conductivity in both MRI and lab sessions. We secured the electrodes on the scalp using self-adhesive tape. We placed the target electrode over the left M1 and the reference electrode over the right orbitofrontal area (see Fig. 1B for a graphical representation of the montage and simulation of the current spread). To achieve this, we used a standard 10–20 system EEG cap to mark the position of C3 and Fp2 and place the active and reference electrodes respectively. Specifically, for anodal stimulation we placed the anode over C3 and the cathode over Fp2 and we reversed this for cathodal sessions. We used the anodal montage in half of the sham sessions, and the cathodal montage in the other half.

In the anodal and cathodal sessions, we stimulated for 20 minutes at 1mA, with 30 seconds ramp-up and 30 seconds ramp-down. Sham stimulation lasted only 30 seconds - also with 30 seconds ramp-up and 30 seconds ramp-down - to give the feeling of active stimulation, in line with well-established protocols used to ensure blinding [36].

### MRI acquisition

We acquired all MRI data on a Siemens MAGNETOM Prisma 3T system, with a 64-channel head coil, at the Centre for Human Brain Health (University of Birmingham). We used the following fMRI parameters: 63 slices, TR = 1620ms, TE = 35ms, matrix size = 84×84×63, voxel size = 2.5×2.5×2.5, no gap, flip angle = 71° and iPAT acceleration factor = 3. We collected 198 volumes for each run of the command-following task.

In addition to the functional scans, we also acquired a high-resolution T1-weighted MPRAGE image, used for anatomical co-registration, using the following parameters: TR = 2000ms, TE = 2.03ms, matrix size = 256×256×208, voxel size = 1×1×1mm, and flip angle = 8°.

We also collected diffusion tensor imaging data, fMRI data during stimulation, and resting state fMRI before and after the stimulation. However, we will analyse and report these in separate papers.

### fMRI paradigm

We used the same paradigm described in [13], whereby we instructed participants to perform a simple thumb movement (adduction-abduction), as fast as possible, in response to an auditory cue (beep) (Fig. 1C and 1D). Participants completed this task before and after tDCS stimulation. The use of a simple task that emulates the command following tasks typically used as part of clinical assessments, enables us to translate our methods to PDOC patients, as well as to study the effects of tDCS on the brain independently of modulations of task performance. We presented the beeps in 8 blocks of 20 seconds, for a total duration of 5 minutes and 20 seconds. Each active block started with the word “move”, and alternated with blocks in which participants were instructed to rest (cued by the word “relax”). Each active block included 7 beeps, which appeared at a variable interstimulus time, ranging from 2 to 3 seconds, to avoid prediction effects. Additionally, we instructed participants to maintain their gaze fixated on a white cross displayed in the centre of a black screen throughout the full duration of the tasks. At the beginning of each run of the task, we displayed the following written instructions: “Start moving your thumb as quickly as you can every time you hear a beep. Stay still when you hear ‘relax’. Make sure you keep looking at the fixation cross at all times”.

### Passive mobilisation

In all sessions (MRI and Lab), we passively moved the participant’s right thumb during the 20 minutes of tDCS stimulation. For this, we used a custom made system that involved a piston attached to the MRI-compatible joystick (Fig. 1D) used to record participants’ responses in the CF-fMRI task. The piston was connected to an air compressor through plastic tubes. We used a Jun-Air Compressor Sj-27 placed in the MRI control room during the MRI sessions, and a Bambi BB24 air compressor (24 litres) in the lab sessions. To open and close the airflow and move the piston, we used an electrically-operated valve, controlled with a Matlab script. In order to achieve a full range of movement (i.e., full abduction-adduction of the thumb), we secured the piston to the participant’s thumb using a rubber band. This set-up was the same for both MRI and lab sessions. We delivered passive movements in blocks of 20 seconds, with 7 movements per block (with a variable time interval of 2-3 seconds), and 20 seconds rest between blocks, to emulate the timings in the command-following task. The total number of passive movements administered during the 20 minutes of stimulation was 210 (i.e., 30 blocks of passive movements and 30 blocks of rest). Importantly, we fitted participants with noise attenuating earphones both in the MRI and the lab sessions, to prevent them from hearing any noise produced by the air-compressor and piston, and being able to anticipate the passive movements as a result (see experimental procedure). Note that the scanner noise provided further masking. We also instructed participants to keep their gaze at a fixation cross throughout. Additionally, in the MRI scanner participants were laying in a supine position and could not see their thumb moving, ensuring no visual feedback.

### fMRI preprocessing and DCM analysis of the command-following task

#### fMRI preprocessing

We used SPM12 on MATLAB 2015b for the preprocessing of the fMRI data and followed a standard pipeline, similar to our earlier study [13]. This included realignment, co-registration between the structural and functional scans, spatial normalisation, and smoothing with an 8mm FWHM Gaussian Kernel.

#### Region selection and time series extraction

DCM uses Bayesian modelling to infer effective connectivity among brain areas from measured brain activity [37]; that is, given a specific network architecture, DCM models how information propagates through different regions to produce the observed data [38]. We followed a similar pipeline to our previous study [13]: First, we identified the canonical pattern of activity on our command-following task at the group level (Fig. 2, green box). To this end, we first built a multi-session 1st-level fixed-effects (FX) model for each individual subject, including all 12 command-following task runs (pre and post stimulation x Day1/5 sessions x 3 testing weeks) and modelling the contrast corresponding to the move vs rest blocks across sessions (non-weighted average). Then, we performed a second-level random-effects (RX) 1-sample t-test on the individual contrasts to assess the group effects. In the resulting map, we identified the group peak of activation for the following motor regions of interest (ROI) obtained from the AAL atlas: left precentral gyrus (M1), left supplementary motor area (SMA), left thalamus (TH), and right cerebellar lobes IV-V and VIII (CB). We thresholded results using a family-wise error (FWE) corrected p<0.05 (see coordinates in bold in Table 1). The resulting group-derived coordinates then acted as a starting point for identifying, in each individual run, the nearest local maxima. We constrained the search of the individual peaks within a sphere centred on the group peak and with a size of 13 mm radius for the left SMA, M1 and right cerebellum, and 7mm for the left thalamus. The difference in the allowed distance from the group coordinates accommodated for the differences in the regions’ anatomical sizes. Individual peaks had to exceed a liberal statistical threshold of uncorrected p<0.05 [37]. When no peak was found for a specific run at p<0.05, we iteratively relaxed the threshold in 0.05 increments until reaching p = 0.25. When no peak could be found at this threshold, we used the group coordinates instead [39]. It is important to mention that we only used these thresholds to identify the individual peak coordinates used for the feature selection (i.e., the extraction of the time series) but not for any statistical analyses. Once we identified the individual peak coordinates for each run, we extracted time series from 4mm radius spherical volumes of interest centred on them.

**Fig. 2.**
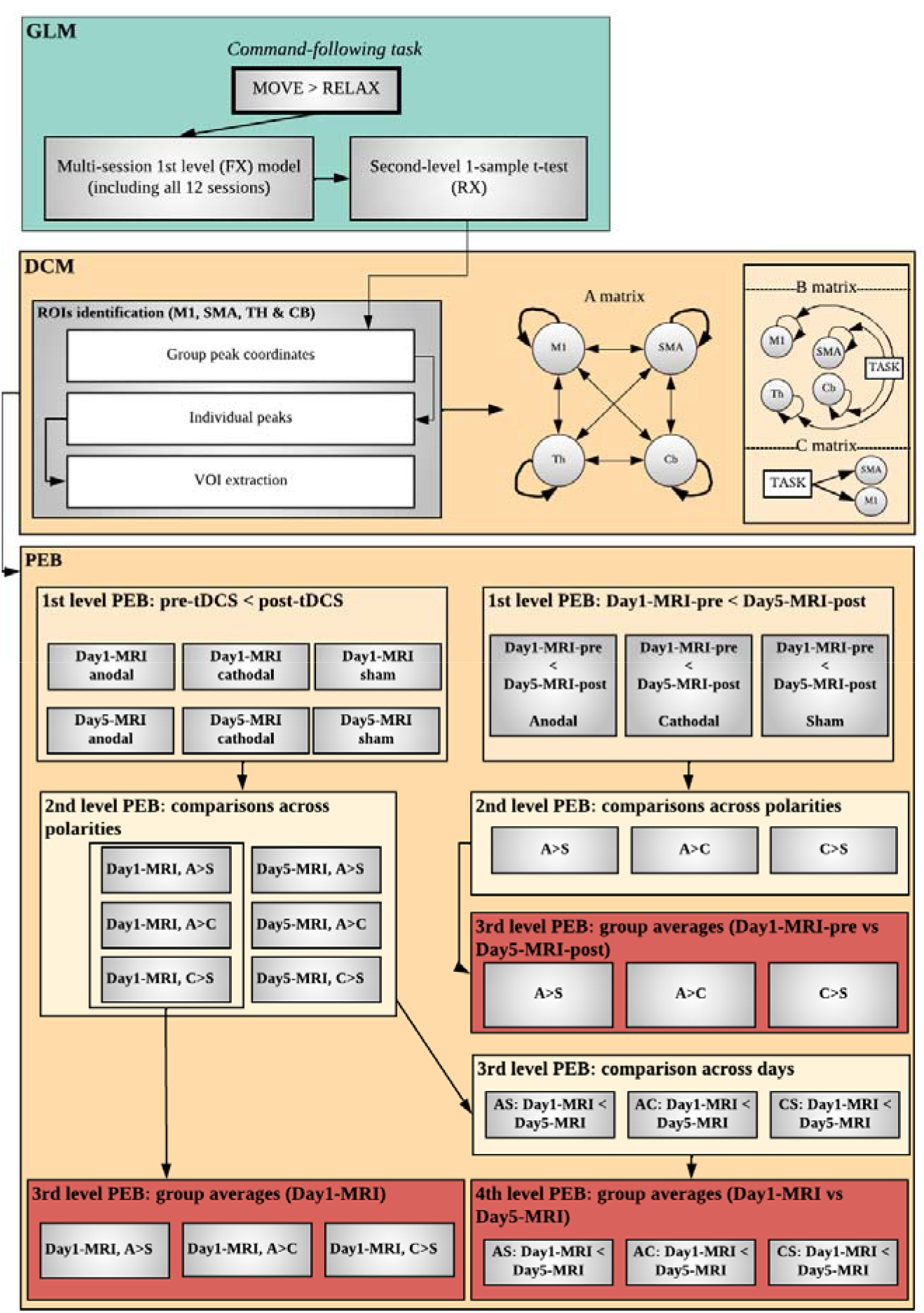
Analysis pipeline. We followed a standard pre-processing protocol (not displayed), followed by general linear model analyses to obtain single-subject and group activation across all runs (*GLM -* panel). We then built and estimated DCMs for each participant and run (see *DCM* panel for a description of our model space). Finally, we used Parametric Empirical Bayes (PEB) to test group differences in effective connectivity (*PEB* panel). Abbreviations: GLM, general linear model; DCM, dynamic causal modelling; FX, fixed-effects; M1, primary motor cortex; SMA, supplementary motor area; TH, thalamus; CB, cerebellum; VOI, volume of interest; PEB, parametric empirical bayes; AS, anodal greater than sham; AC, anodal greater than cathodal; CS, cathodal greater than sham; A, anodal; C, cathodal; S, sham.

**Table 1.** Canonical activation during command-following.

| **Region** | **Cluster P** | **Cluster size** | **Peak P** |  | **F** | **MNI coordinates** |
| --- | --- | --- | --- | --- | --- | --- |
|  | *FWE-corrected* | *in mm3* | *Peak FWE-corrected* | *uncorrected* |  |  |
| **Cerebellum** | <0.001 | 3969 | <0.001 | <0.001 | 11.871 | **[12;-46;-19]** |
|  |  |  | <0.001 | <0.001 | 11.337 | [9;-52;-13] |
|  | <0.001 | 729 | <0.001 | <0.001 | 9.465 | [12;-58;-43] |
|  | <0.001 | 162 | 0.005 | <0.001 | 7.399 | [27;-49;-46] |
| **M1** | <0.001 | 5697 | <0.001 | <0.001 | 10.570 | **[-36;-19;56]** |
|  |  |  | <0.001 | <0.001 | 9.695 | [-27;-19;74] |
|  |  |  | <0.001 | <0.001 | 9.174 | [-33;-13;65] |
|  | 0.026 | 27 | 0.008 | <0.001 | 7.126 | [-42;-1;20] |
| **SMA** | <0.001 | 3834 | <0.001 | <0.001 | 9.812 | **[-6;-1;56]** |
|  |  |  | 0.004 | <0.001 | 7.450 | [-9;8;44] |
| **Thalamus** | 0.002 | 297 | <0.001 | <0.001 | 9.506 | **[-18;-22;11]** |

Results from the random effect group analyses (RFX, 1-sample t-test) for the command-following task, including all 12 MRI sessions per participant. Results survived a threshold of FWE-corrected $p < 0.05$ . We used the peak coordinates for each region (highlighted in bold) as a starting point to extract the time series for the DCM analyses.
Abbreviations: FWE, family wise error; M1, primary motor cortex; SMA, supplementary motor area.

#### Individual level DCM specification and definition of model space

After the extraction of the time series, we specified dynamic causal models (Fig.2, DCM panel) at the individual level with a deterministic model for BOLD signal and the following parameters: one state per region, bilinear modulatory effects, and mean-centred inputs. DCM requires the specification of 3 matrices modelling the connectivity within- and between-regions (A matrix), the modulations of effective connectivity due to experimental conditions (B matrix), and driving inputs (C matrix), which briefly ‘ping’ specific regions in the system at the onset of each block [37].

We defined our model space using the same parameters to those in our earlier study [13]: a fully connected A matrix (i.e., a model in which all self- and between region connections are switched on) including left M1, left SMA, left thalamus and right cerebellum, a B matrix including the effect of the motor task as modulatory input on all self-connections, and a C matrix modelling driving inputs to cortical regions only (M1 and SMA).

#### tDCS effects on effective connectivity

With the above DCMs, we employed Parametric Empirical Bayes (PEB) to remove the parameters that were not contributing to the model evidence, and to evaluate group effects and between subject variability [39]. In order to test the effects of tDCS combined with passive mobilisation on the connections in our model, we ran three separate hierarchical PEB analyses [39], starting from the 12 neural models (DCMs) estimated per participant (i.e., one per CF-fMRI run).

1. To assess the effects of a single tDCS session, we first built 3 *within-subject* PEBs (1^st^-level) per participant encoding the differences between pre- and post-stimulation (pre-tDCS < post-tDCS) for each polarity on the first MRI session. We then entered these into 3 *within-subject* PEBs (2^nd^ level) for each participant that encoded comparisons between polarities (Anodal > Sham, Anodal > Cathodal, Cathodal > Sham) on the differences between pre- and post tDCS. Finally, we specified 3 group (*between-subject*) PEBs encoding the commonalities across participants for each 2^nd^-level PEB, and included sex, age, and handedness score as regressors of non-interest (3^rd^ level). The 3 final PEBs therefore encode increases (i.e., pre-tDCS<post-tDCS) in effective connectivity that are greater after anodal tDCS as compared to sham (i.e., anodal > sham), after anodal tDCS as compared to cathodal tDCS, or after cathodal tDCS as compared to sham respectively.
2. To assess the cumulative effects of 5 tDCS sessions, we replicated the above steps but encoding the differences between the pre-tDCS run in Day1-MRI and the post-tDCS run in Day5-MRI for the 1^st^ level of the hierarchy. 2^nd^ and 3^rd^ level PEBs were as above.
3. Finally, to establish whether the responsiveness to tDCS increases after multiple sessions, we replicated the 1^st^ and 2^nd^ level steps in (1) above for the data on Day5-MRI. Then created a further *within-subject* PEB modelling the differences between sessions (Day1 < Day5) for each individual participant (3^rd^ level). Finally, a between subjects PEB modelled group effects (mean across participants), and included sex, age, and handedness score as regressors of no interest (4^th^ level).

Finally, we used Bayesian Inference to invert the model for each subject and obtain an estimate of the parameters that best explain the data, while minimising complexity. First, we applied Bayesian Model Reduction (BMR) on the final 3 PEBs above (i.e., those encoding group commonalities), to reduce connections that were not adding to the model evidence. Finally, we used Bayesian Model Average (BMA) to estimate the average parameters across participants for each connection that remained switched on. We used a threshold of a posterior probability > 95%, as we have previously done [13].

In addition, to assess the stability of our parameters across sessions, we created 3 PEBS modelling the average across sessions on the baseline pre-tDCS run on Day 1 for each polarity (i.e., run unaffected by any stimulation or passive mobilisation) and used BMR and BMA to estimate parameters as above.

### Motion tracking

We analysed the motion data for the CF-fMRI task with a custom MATLAB 2017b script, as we did in our previous study [13]. We first low-pass filtered the data at 15Hz and calculated the Euclidean distance of the x-y position. To identify the onset and the end of each movement in the arrays containing our motion tracking data, we used the MATLAB function *findchangepts*, which can detect abrupt signal changes given a vector x with N elements and returns the index at which the mean of x changed most significantly. When more than two changes were identified, we used the first and last ones to determine the beginning and end of the movement. We excluded movements where no change was detected, which was usually due to participants not responding to that specific stimulus (on average we removed <1 movement per session, maximum of 12). For each movement we calculated velocity and acceleration at each timepoint, and then derived the mean velocity and peak acceleration. We then averaged those session means to give the mean velocity and peak acceleration for the whole run. We also derived reaction times, defined as the time interval between the auditory stimulus and onset of the movement, identified as per above.

We calculated the means of those values across each subject/run and excluded measurement errors, defined as values with a z-score > 3.0, that were present due to the joystick being incorrectly calibrated. Out of 264 runs (per analysis), we removed 14 from the reaction times analysis, 1 from the mean velocity analysis and 36 from the peak acceleration analysis. Finally, for each of the three metrics (i.e., reaction time, mean velocity and peak acceleration) we computed three Bayesian repeated measures ANOVAs on JASP, v. 0.16.3 [40] to emulate the DCM analyses above:

1. A Polarity (anodal, cathodal and sham) x Time (pre- vs post-tDCS) ANOVA on the data from Day1-MRI tested the effects of a single tDCS session.
2. A Polarity x Multiple session time (Day1-MRI pre-tDCS vs Day5-MRI post-tDCS) ANOVA tested the effects of 5 sessions.
3. Polarity x Time x Day (Day1-MRI vs Day5-MRI) ANOVA tested whether within-session responsiveness to tDCS is affected by the number of sessions.

For each ANOVA, we calculated the evidence in the data for including the interaction (e.g., polarity x time) as a predictor (BF_incl_); that is, whether the models including the interaction explain the data significantly better than those without it [41]. We interpreted BF_incl_ > 3 as substantial evidence in the data for including the interaction, BF_incl_ > 10 as strong evidence, and BF_incl_ > 100 as very strong evidence, as per standard guidelines on interpreting Bayes factors [42]. Importantly, for BF_incl_ < 3, we reported the exclusion Bayes factor (BF_excl_) instead, which quantifies the opposite effect (the evidence for models excluding the interaction as compared to those including it).

### Blinding

To test the success of our blinding protocol, we used McNemar’s test on participants’ responses as to whether they believed they had / had not received real tDCS on days 1 and 5 (MRI sessions).

## Results

### Task activation across sessions

The RFX second-level analysis of brain activation during the active movements to command across sessions showed statistically significant clusters (FWE-corrected p<0.05) in all of four ROIs considered in our analyses (see Table 1 and Fig. 3).

**Fig. 3.**
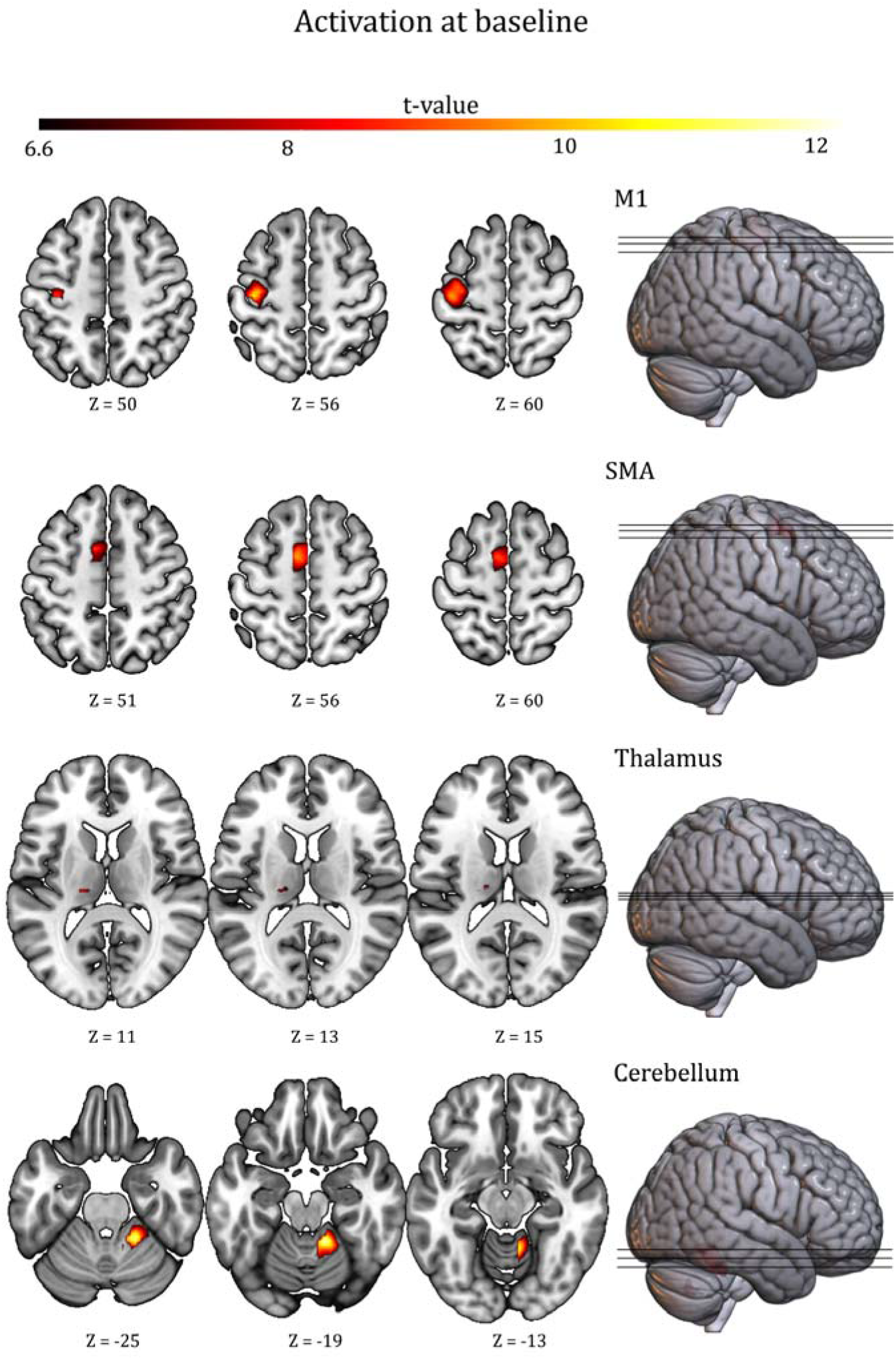
Activation at baseline. Brain activation at the group level (RFX, 1-sample t-test) for the command-following task. The general linear model included differences between ‘move’ and ‘rest’ blocks on a multi-session 1st level (FFX) model for each subject - using all 12 MRI runs. The activation maps are shown at p<0.05 FWE-corrected for M1, SMA, cerebellum and thalamus ROIs, and rendered on a standard template (152 template in MRIcroGL). Z indicates the Montreal Neurological Institute z coordinate.

**Fig. 4.**
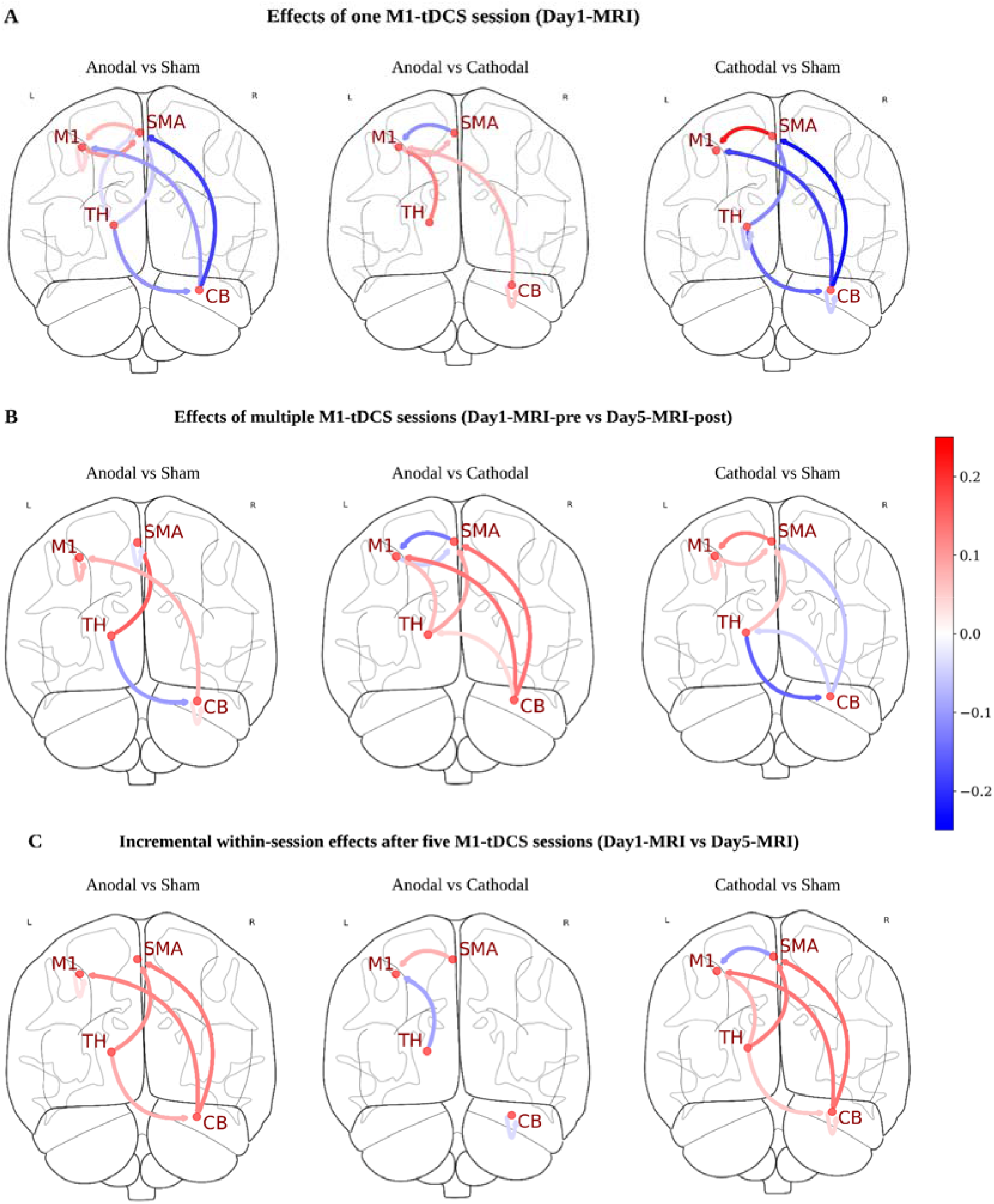
Effects of M1-tDCS on neural dynamics. Results from our Parametric Empirical Bayes (PEB) analyses on Day1-MRI (**A**), Day1-MRI-pre vs Day5-MRI-post (**B**) and pre- vs post- on Day1-MRI vs Day5-MRI (C). Blue lines indicate decreases and red lines indicate increases in connectivity strength. Notice that we converted self-connections from their original unitless log-scale values to Hertz and added 0.5 to obtain the direction and magnitude of the self-connectivity change and display it in a way that is comparable with between region connections.

### Effects of M1-tDCS on motor-network dynamics

#### Effects of one M1-tDCS session (Day1-MRI)

A single anodal M1-tDCS session (Fig. 5, panel A) increased excitation from M1 to SMA, when compared to both cathodal- and sham-tDCS, and from thalamus to M1 as compared to cathodal tDCS only. Additionally, when compared to sham only, anodal tDCS decreased M1 self-inhibition as well as excitation from SMA to thalamus. In turn, a single session of cathodal tDCS, increased cerebellar self-inhibition, when compared to both anodal and sham, and thalamic self-inhibition when compared to sham only. Both anodal and cathodal tDCS increased excitation from SMA to M1 and inhibition from cerebellum to M1, although these changes were more pronounced for cathodal and anodal tDCS respectively. Similarly, both polarities increased inhibition from thalamus to SMA and cerebellum, as well as from cerebellum to SMA, with no differences between polarities in this case. In terms of task modulations on each self-connection (matrix B), we found no significant effects for any polarity.

**Fig. 5.**
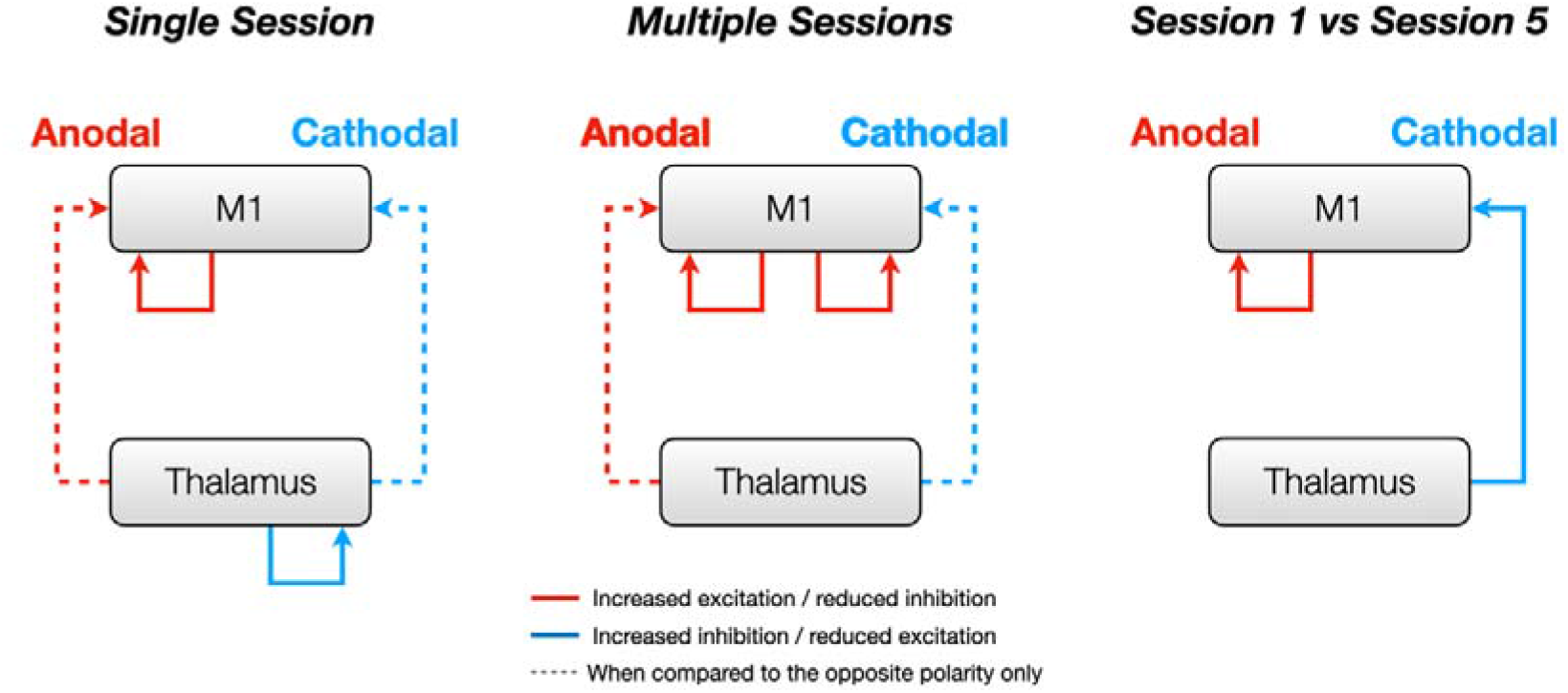
Effects of M1-tDCS on thalamo-M1 coupling. Summary results for the thalamus-M1 complex after 1 session (left), 5 sessions (middle), and when comparing within-session changes between day 1 and day 5 (right). Dashed lines represent effects that were only significant in the comparison between polarities (but not when each polarity was compared to sham).

See Fig. S1 for baseline (pre-tDCS) connectivity values.

#### Effects of multiple M1-tDCS sessions (Day1-MRI-pre vs Day5-MRI-post)

After 5 sessions, anodal tDCS increased excitation from thalamus to M1 when compared to cathodal tDCS only. It also increased self-inhibition in SMA and reduced self-inhibition in the cerebellum as compared to sham only. Finally, it increased excitation from cerebellum to M1 when compared to both cathodal and sham. Both anodal- and cathodal tDCS increased excitation from thalamus to SMA, although this was significantly stronger for anodal stimulation. Similarly, both polarities reduced M1 self-inhibition and excitation from thalamus to cerebellum compared to sham, with no significant difference between polarities. In turn, cathodal tDCS increased excitation between M1 and SMA (in both directions) and inhibition from cerebellum to thalamus and SMA, both as compared to anodal tDCS and sham. As above, we found no significant effects in task modulations for either polarity.

#### Incremental within-session effects after five M1-tDCS sessions (Day1-MRI vs Day5-MRI)

Compared to the first session, the fifth session of anodal M1-tDCS (Fig. 5, panel B) reduced M1 self-inhibition as compared to sham only. In addition, both polarities increased excitation from thalamus to SMA and from cerebellum to M1, and reduced inhibition from thalamus to cerebellum and cerebellum to SMA, with no differences across polarities.

In turn, the fifth session of cathodal tDCS reduced thalamo-M1 inhibition, and decreased cerebellar self-inhibition and excitation from SMA to M1. All of these were polarity specific, i.e., significant both when compared to anodal tDCS and sham. As above, we found no significant effects in task modulations for either polarity.

### Effects of M1-tDCS on behavioural metrics

We found substantial evidence in support of the lack of an interaction (BF_excl_ > 3), for reaction times and mean velocity on Day1-MRI and Day1-MRI pre vs Day5-MRI post, as well as for peak acceleration on Day1-MRI only. In addition, we found anecdotal evidence (i.e., BF_excl_ =1-3) for the lack of an interaction on mean velocity for Day1-MRI vs Day5-MRI and peak acceleration for the comparisons Day1-MRI pre vs Day5-MRI post and Day1-MRI vs Day5-MRI. We report means and standard deviations for each condition in Table S1 along with the BF_excl_ values and their respective prior and posterior inclusion probability, for all ANOVAS performed. Single-subject values are in Fig. S2.

### Efficacy of blinding protocol

We found no significant differences in the number of times that participants perceived sham and anodal/cathodal stimulation as real stimulation for either Day1-MRI (χ^2^=9, p=0.82) or Day1-MRI (χ^2^=11, p=0.20)

## Discussion

This study focused on characterising the effects of tDCS paired with passive mobilisation on thalamo-cortical coupling during a motor command-following task; a circuit with an established mechanistic role in the behavioural responsiveness in PDOC [17]. We have previously shown that one anodal M1-tDCS session administered at rest can reduce thalamic self-inhibition during command following. However, against prediction, these changes were not followed by increased excitation towards M1 [13]. Here, we demonstrated that, when paired with passive mobilisation, one session of anodal M1-tDCS can indeed increase thalamus to M1 excitation, (and reduce M1 self-inhibition as a result), during the same command following task. This suggests that pairing passive mobilisation with stimulation could have greater therapeutic effects in PDOC, particularly in those patients whose diminished responsiveness is linked to reduced thalamo-cortical connectivity [17,18]. We have previously shown that both an excitatory engagement of the thalamus and thalamus to M1 coupling are necessary for the execution of voluntary motor responses to command in PDOC [17]. It is also well known that damage to the thalamus and / or its cortical projections are common in PDOC, and that these abnormalities correlate with the level of reduced responsiveness in these patients [17,18,43,44]. As individual patterns of damage are highly heterogeneous across PDOC patients, one could speculate that, depending on whether the primary damage affects mostly the thalamus or its connections to M1, different patients may benefit most from stimulation being delivered at rest [13] or combined with passive mobilisation (as reported here) respectively. Further studies in PDOC patients themselves are required to ascertain this.

Leaving implications for clinical improvements aside momentarily, our previous study [13] revealed unexpected effects after one session of cathodal M1-tDCS, which, against predictions, led to an increase in excitation from thalamus to M1. We attributed this to a compensatory mechanism to overcome the strong inhibition in M1 that followed this polarity and keep performance at a baseline level. In contrast, here we found that anodal and cathodal stimulation showed opposing effects over this connection (i.e., excitatory for anodal and inhibitory for cathodal), in line with our predictions and existing literature. Pairing tDCS with a relevant task is indeed regarded as best practice [45]. In fact, there is much evidence that the effect of tDCS is greatly dependent on the neural state during stimulation, and that using a task that engages the relevant neural circuit can increase the spatial precision of tDCS on the target networks [22]. This can be of even greater importance in patients like those in PDOC, where gross brain abnormalities interfere with an accurate identification of the targets at the scalp level[1]. Despite this, to our knowledge, all previous tDCS studies in PDOC administered the stimulation at rest[1]. While this is unsurprising, due to the patients’ inability to engage in an active task, we here provide an alternative passive task (for protocols targeting motor areas) that is suitable in this cohort.

Importantly, we know that multiple sessions of tDCS can lead to more robust clinical improvements in PDOC (see [1] for a review). This is indeed a well-documented phenomenon in therapeutic tDCS applications across clinical conditions[46]. Our study offers a window into the neural underpinnings of the cumulative effects of multi-session tDCS, and revealed a somewhat complex picture, particularly for cathodal stimulation. First, after 5 tDCS sessions, anodal stimulation continued to increase thalamus to M1 excitation and to decrease M1 inhibition as compared to baseline. Similarly, cathodal tDCS decreased thalamus to M1 excitation both after 1 and 5 sessions (although significantly less so after 5). However, after 5 sessions, cathodal tDCS also decreased M1 inhibition, and did so at a similar level to anodal tDCS. Moreover, after 5 sessions, cathodal tDCS lost the influence on increasing thalamic self-inhibition that it had after 1. Crucially, the baseline (pre-tDCS) connectivity within and between these two regions was very stable across polarity sessions (see figure S1). There were other effects outside these key areas of interest (thalamus, M1 and their connection) that only became apparent or changed tone after 5 sessions of either polarity. For example, the thalamus changed from inhibiting SMA after one session of either anodal or cathodal tDCS to exciting it after 5 sessions of either polarity. In turn, 5 sessions of anodal tDCS were required to reduce cerebellar self-inhibition, while the effects of cathodal tDCS over the cerebellum in session 1 disappeared after 5 sessions. Looking at the baseline connectivity, it becomes clear that the existing neural state modulates the effect of each polarity to bring the network to an optimal level that maintains behavioural performance. Overall, our results suggest that future research should not necessarily expect opposing effects for the two polarities across the network of interest, nor linear, incremental effects after multiple sessions. The comparison of within-session effects sheds further light into this (Figure 5, panel C). For example, anodal tDCS had had a stronger effect on decreasing M1 self-inhibition on the fifth session as compared to the first, suggesting a potentiation of the events (rather than a simple accumulation). In contrast, the increased excitation from thalamus to M1 related to this polarity did not exhibit significant within-session differences (i.e., it was similar in sessions 1 and 5). Finally, the inhibitory effect of cathodal tDCS over this connection was smaller in the fifth session as compared to the first.

Most of our knowledge about tDCS comes from studies focusing on local effects over M1, and much less is understood about how the modulations propagate across other regions in the motor network. It is thus not surprising that the local changes on self-inhibition we identified here are consistent with prior literature, while long-range changes appeared more complex. There is now substantial evidence that anodal tDCS over M1 increases cortical excitability [47,48], and this modulation of neural activity is associated with changes in synaptic plasticity [49], including early- (e-LTP) and late long-term potentiation (l-LTP) when an appropriate task is used. Crucially, l-LTP requires repeated sessions of stimulation and can lead to structural plasticity that may contribute to more robust tDCS aftereffects[50]. Furthermore, both anodal and cathodal tDCS have a similar impact on synaptic plasticity[50], consistent with our changes after 5 sessions showing reduced M1 self-inhibition for both polarities (albeit stronger for anodal). While it is possible that e-LTP and l-LTP may be one of the mechanisms explaining the local effects on M1 in our current study, especially as we did not find an effect of anodal tDCS on M1 self-connectivity when delivering tDCS at rest in our previous study[13], the relationship between synaptic mechanisms and effective connectivity metrics is not well understood. In fact, DCM does not model the activity of individual neurons, which are characterised by fast dynamics, but slower dynamics - in the scale of seconds - that arise from the synergy of several neuronal populations instead [37]. Therefore, the comparison between DCM results and neuronal models is not straightforward. Within the DCM framework, a reduction in self-inhibition reflects a lowered rate of decay of neuronal activity, which in turn indicates increased susceptibility to afferent inputs into that region[37]. In our case, the reported reduction in M1 self-inhibition would make M1 more sensitive to the other nodes in the network (including the excitation coming from thalamus). This way, even when the increase in excitation from thalamus to M1 seemed to be maintained (rather than enhanced) across the sessions, the enhanced responsiveness in M1 to this thalamic excitation after 5 sessions would predict a stronger clinical effect. Importantly, within our network, M1 represents the final motor output for the control of muscles, due to its projections to the spinal cord [51]. As such, increasing M1 excitability via a combination of reduced M1 self-inhibition and increased thalamo-M1 excitation might successfully facilitate the initiation of movements in those PDOC patients where the cortico-thalamic tracts are partially spared.

It is worth highlighting here that we designed our command-following task to emulate those used in clinical settings to evaluate residual awareness in PDOC [15] and, as a result, our task is insensitive to tDCS-related modulations of behaviour in healthy participants. Indeed, consistently with our previous study [13], we found no effect of tDCS on behavioural measures in our cohort. Therefore, further studies in PDOC are required to confirm whether the neural effects reported here indeed translate into clinically meaningful changes in patients. If successful, an increase in thalamo-cortical excitation in PDOC could improve patients’ behavioural responsiveness and impact their quality of life by facilitating engagement with rehabilitation as a result [52]. Additionally, an improvement in responsiveness would also decrease misdiagnosis levels, especially in those patients with CMD, by uncovering patients’ real level of awareness.

## Conclusions

Overall, we showed that anodal tDCS coupled with passive mobilisation of the thumb has short- and long-range effects on motor-network dynamics during command-following. Specifically, it can reduce M1 self-inhibition and increase excitation from thalamus to M1. These effects are enhanced after 5 sessions of stimulation. While we tested our protocol on healthy participants, we designed it to be easily integrated into PDOC care as part of routine physical therapy.

## Credit authorship contribution statement

Davide Aloi: Investigation, Data curation, Formal analysis, Methodology, Writing – original draft, Writing – review & editing. Roya Jalali: Investigation, Methodology, Writing – review & editing. Sara Calzolari: Methodology, Writing – review & editing. Melanie Lafanechere: Investigation, Writing - review & editing. R. Chris Miall: Methodology, Writing – review & editing. Davinia Fernández-Espejo: Visualization, Funding acquisition, Data curation, Methodology, Supervision, Writing – original draft, Writing – review & editing.

## Acknowledgements

This work was supported by generous funding from the Medical Research Council (MR/P02596X/1; DF-E). DA was supported by a scholarship from The Wellington Hospital and the University of Birmingham.

We would also like to thank David McIntyre and Stephen Allen for their help with the construction of the passive mobilisation device.

## Supplementary Material

### Supplementary Methods

#### C-Matrix definition

As we have previously done [13], to establish the set of driving inputs (C matrix) that best fitted our data, we first created DCM models that included driving inputs to all 4 regions (12 DCM models per participant, one per fMRI run). We then created a second level within-subject PEB per participant, to model the commonalities between all 12 runs. We subsequently entered the resulting 22 PEBs into a third level PEB, to model the similarities across all participants. We included in the model a constant encoding the group mean, as well as the following nuisance regressors: sex, age, and Edinburgh Handedness Inventory score (all mean-centred). To do this, we applied Bayesian Model Reduction (BMR) to iterate over the reduced models, and then used Bayesian Model Average (BMA) to determine the average connectivity parameters[37]. We used a 95% posterior probability threshold for free energy (i.e., comparing the evidence for all models where a particular connection/input is included versus when it is excluded). Like in our previous study[13], this step showed robust support (>95% posterior probability) for the inclusion of driving inputs to cortical regions only (M1 and SMA). We therefore modelled the final DCMs for all participants with all self- and between-region connections (A matrix), modulatory inputs to each self-connection (B matrix) and driving inputs to M1 and SMA (C matrix).

## Supplementary Tables and Figures

**Table S1.** Effects of tDCS on behavioural metrics (V2)

| | | | | | Analysis of Effects: $BF_{\text{excl}}$ ($P(\text{excl}) \rightarrow P(\text{excl}\vert \text{data})$) | | |
| --- | --- | --- | --- | --- | --- | --- | --- |
| Metrics | Day | Polarity | Baseline | Post-tDCS | Day1-MRI only | Day1-MRI pre vs Day5-MRI post | Day1-MRI vs Day5-MRI |
| **Reaction times (s)** | Day1-MRI | Anodal | 0.221 (± 0.032) | 0.220 (± 0.044) | 4.732 (0.200 → 0.082) | 7.689 (0.200 → 0.106) | 0.506 (0.053 → < 0.000) |
| | | Cathodal | 0.244 (± 0.058) | 0.230 (± 0.057) | | | |
| | | Sham | 0.225 (± 0.051) | 0.233 (± 0.056) | | | |
| | Day5-MRI | Anodal | 0.220 (± 0.040) | 0.214 (± 0.047) | | | |
| | | Cathodal | 0.231 (± 0.056) | 0.230 (± 0.049) | | | |
| | | Sham | 0.217 (± 0.037) | 0.214 (± 0.040) | | | |
| **Mean velocity (cm/s)** | Day1-MRI | Anodal | 8.801 (± 5.053) | 7.242 (± 3.741) | 5.323 (0.200 → 0.090) | 3.936 (0.007 → 0.026) | 2.227 (0.053 → < 0.000) |
| | | Cathodal | 8.910 (± 4.270) | 8.196 (± 3.691) | | | |
| | | Sham | 7.924 (± 4.176) | 7.881 (± 4.489) | | | |
| | Day5-MRI | Anodal | 8.463 (± 4.042) | 7.880 (± 4.203) | | | |
| | | Cathodal | 8.196 (± 3.828) | 9.100 (± 4.367) | | | |
| | | Sham | 8.513 (± 3.756) | 8.731 (± 4.573) | | | |
| **Peak acceleration ($m/s^2$)** | Day1-MRI | Anodal | 39.495 (± 24.847) | 32.871 (± 15.274) | 4.143 (0.200 → 0.076) | 1.531 (0.025 → 0.038) | 1.066 (0.053 → < 0.002) |
| | | Cathodal | 38.049 (± 22.992) | 37.162 (± 19.804) | | | |
| | | Sham | 34.160 (± 17.280) | 37.660 (± 21.256) | | | |
| | Day5-MRI | Anodal | 38.837 (± 21.750) | 46.071 (± 33.901) | | | |
| | | Cathodal | 42.245 (± 22.053) | 46.245 (± 30.373) | | | |
| | | Sham | 35.989 (± 16.956) | 29.900 (± 10.887) | | | |

Statistics for the Bayesian repeated measures ANOVAs (Day1-MRI only, Day1-MRI pre vs Day5-MRI post and Day1-MRI vs Day5-MRI), on behavioural metrics. We report exclusion Bayes factors for the interaction effect between the considered factors for each Bayesian ANOVA. Abbreviation: $BF_{\text{excl}}$ , Bayesian evidence for the exclusion of the interaction in the model, against the matched model; $P(\text{excl})$ , prior inclusion probability; $P(\text{excl}|\text{data})$ , inclusion probability; cm, centimetres; s, seconds; m, metre.

**Fig. S1.**
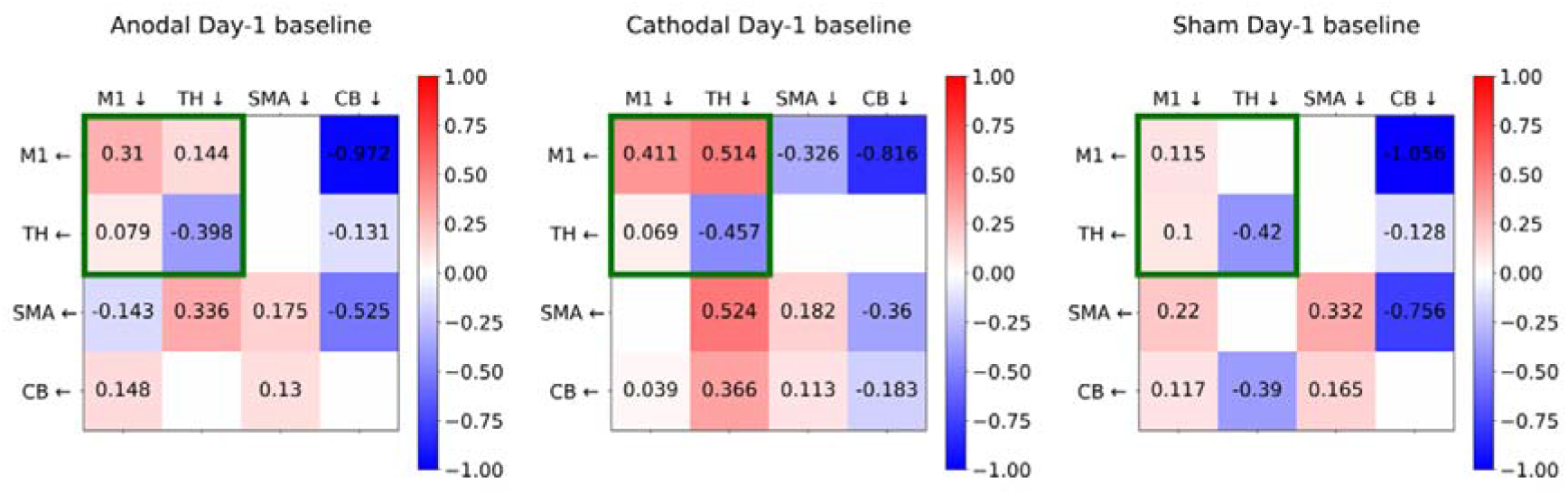
Effective connectivity on the pre-tDCS run on Day1-MRI session. Group average of effective connectivity for the pre-tDCS run on the first session for anodal (left), cahotodal (middle), and sham (right) sessions. Arrows represent the direction of the connectivity in each cell. For between-region connections red cells indicate excitatory coupling and blue cells indicate inhibitory coupling. Self-connections are always inhibitory and red/blue cells indicate increased/decreased inhibition respectively. Green boxes highlight the key connections of interest. We display connections that survive a 95% posterior probability threshold for free energy.

**Fig. S2.**
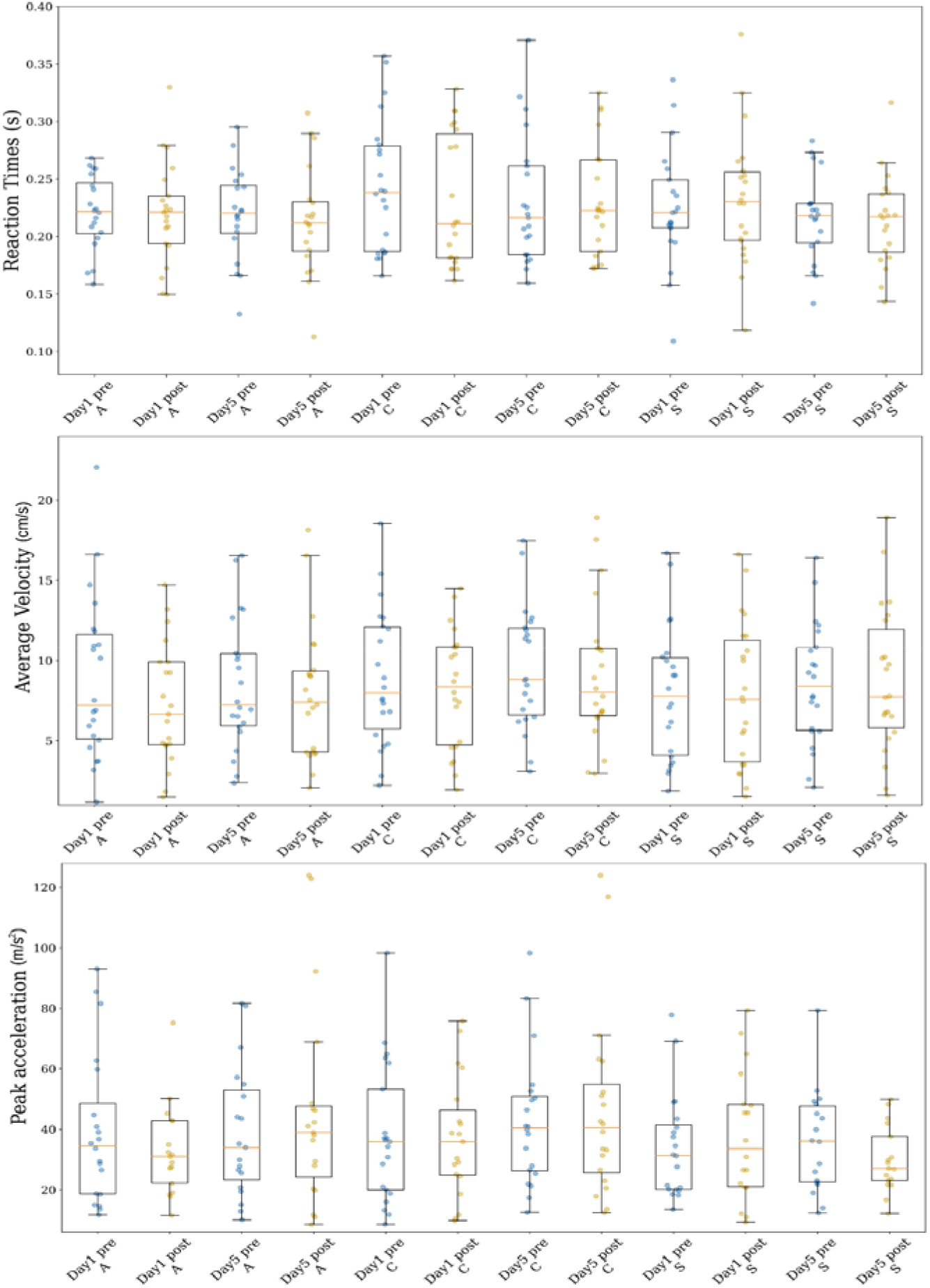
Effects of M1-tDCS on behavioural metrics. Effects of M1-tDCS on behavioural command following. Scatter plots displaying the average values per condition for the reaction times (top figure), average velocity (centre) and peak acceleration (bottom). In each column, each dot represents the averaged value for one participant. Data-points corresponding to measurement errors (defined as values with a Z-score > 3.0) are not displayed. Abbreviations: Day1, first day of the testing week (i.e., Day1-MRI); Day5, last day of the testing week (i.e., Day5-MRI); A, anodal stimulation; C, cathodal stimulation; S, sham stimulation; pre, before stimulation; post, after stimulation.

## Notes

### Competing Interest Statement

The authors have declared no competing interest.

